# Refining the composition of the *Arabidopsis thaliana* 80S cytosolic ribosome

**DOI:** 10.1101/764316

**Authors:** Karzan Jalal Salih, Owen Duncan, Lei Li, Josua Troesch, A. Harvey Millar

**Author notes:** Correspondence: Prof. Harvey Millar.

## Abstract

The cytosolic 80S ribosome is composed of protein and RNA molecules and its function in protein synthesis is modulated through interaction with other cytosolic components. Defining the role of each of the proteins associated with ribosomes in plants is a major challenge which is hampered by difficulties in attribution of different proteins to roles in ribosome biogenesis, the mature cytosolic ribosome (r-proteins) or to the broader translatome associated with functioning ribosomes. Here we refined the core r-protein composition in plants by determining the abundance of proteins in low, partially and highly purified ribosomal samples from *Arabidopsis thaliana* cell cultures. To characterise this list of proteins further we determined their degradation (K_D_) and synthesis (K_S_) rate by progressive labelling with ^15^N combined with peptide mass spectrometry analysis. The turnover rates of 55 r-proteins, including 26 r-proteins from the 40S subunit and 29 r-proteins from the 60S subunit could be determined. Overall, ribosome proteins showed very similar K_D_ and K_S_ rates suggesting that half of the ribosome population is replaced every 3-4 days. Three proteins showed significantly shorter half-lives; ribosomal protein P0D (RPP0D) with a half-life of 0.5 days and RACK1b and c with half-lives of 1-1.4 days. The ribosomal RPP0D protein is a homolog of the human Mrt4 protein, a trans-acting factor in the assembly of the pre-60S particle, while RACK1 has known regulatory roles in cell function beyond its role as a 40S subunit. Our experiments also identified 58 proteins that are not from r-protein families but co-purify with ribosomes and co-express with r-proteins in Arabidopsis. Of this set, 26 were enriched more than 10-fold during ribosome purification. A number have known roles in translation or ribosome-association while others are newly proposed ribosome-associated factors in plants. This analysis provides a more robust understanding of Arabidopsis ribosome content, shows that most r-proteins turnover in unison *in vivo*, identifies a novel set of potential plant translatome components, and reveals how protein turnover can identify r-proteins involved in ribosome biogenesis or regulation in plants. Data are available via ProteomeXchange with identifier PXD012839.

## Introduction

The ribosome uses information encoded by messenger RNA (mRNA) to drive peptide bond formation resulting in protein synthesis (Thompson et al., 2016). All ribosomes consist of two subunits, a large and small, and each contain ribosomal RNAs (rRNAs) and ribosomal proteins (r-proteins). The 80S eukaryotic ribosome is larger and more complicated than the 70S prokaryotic ribosome due to the presence of additional r-proteins and rRNAs (Ben-Shem et al., 2010). The composition, structure and organization of r-proteins and their distribution between small and large subunits have been studied in detail in bacteria (*E.coli*) (Schuwirth et al., 2005) yeast (*S. cerevisiae*) (Jenner et al., 2012), *Tetrahymena thermophile* (Klinge et al., 2011), humans (*Homo sapiens*) (Khatter et al., 2015) and plants (*Arabidopsis thaliana)* (Barakat et al., 2001; Carroll et al., 2008; Hummel et al., 2015).

Based on gene annotation, the 80S cytosolic ribosome in *Arabidopsis thaliana* is understood to contain 81 different r-proteins and four ribosomal RNAs, including 33 proteins in the small subunit and 48 in the large subunit (Barakat et al., 2001). There is a high degree of conservation of ribosomal proteins between *Arabidopsis thaliana* and mammals with the exception of the acidic phosphoprotein (RPP) family, also known as acidic stalk P3 (RPP3), which only exists in plants (Carroll et al., 2008). Many of the Arabidopsis r-proteins are encoded by multiple genes. In a recent survey, 102 Arabidopsis genes were predicted to encode the 33 r-proteins of the small subunit and 146 genes to encode the 48 r-proteins of the large subunit; on average there are 3 predicted Arabidopsis genes for each r-protein type (Carroll et al., 2008). However, in experimental assessments of r-protein content in isolated Arabidopsis ribosomes there are varying sets of r-proteins reported, indicating technical issues with different approaches or potentially variant ribosome contents from different Arabidopsis tissue sources (Chang et al., 2005; Giavalisco et al., 2005; Carroll et al., 2008; Carroll, 2013; Hummel et al., 2015). Each of these reports also identified proteins in ribosome preparations that were not annotated as members of r-protein families. However, each study had limited experimental strategies to determine if such proteins were simply contaminants of ribosome preparations in the different plant tissues used or if they were real ribosome-associated proteins *in vivo*.

Here we refined the core ribosome protein composition by determining the enrichment of r-proteins during purification of ribosomes from *Arabidopsis thaliana* cell cultures. This protein list was further refined by determining the co-degradation and co-synthesis rates of r-proteins during cell growth. We identified a number of proteins that are not from r-protein families but clearly associate and co-purify with Arabidopsis cytosolic ribosomes. This analysis has provide a more robust understanding of Arabidopsis ribosome content, multiple lines of evidence to justify protein inclusion in the core cytosolic 80S r-protein set, a novel set of potential translatome components and evidence that some ribosomal proteins turnover much more rapidly that the ribosome as a whole, indicating potential roles for these proteins in ribosome biogenesis and/or regulation.

## Results

### Enrichment proteomics of 80S cytosolic r-proteins

To quantify the enrichment of proteins during the process of ribosome purification, ribosomal particles from 7 day old *Arabidopsis thaliana* cell culture were extracted, subjected to a three-step enrichment process and analysed in triplicate. Low purity, partially-enriched and highly-enriched ribosome samples were obtained by differential and density-gradient centrifugation as outlined in methods. Proteins in each sample were digested with trypsin and subjected to data-dependent (shotgun) analysis by reversed-phase LC peptide mass spectrometery. The total number of non-redundant proteins identified in all 3 replicates decreased 3-fold as the level of enrichment increased **(Figure 1A**), while the number of proteins annotated as 80S cytosolic r-proteins increased step by step through the purification **(Figure 1B**). Details of the identified proteins are provided in **Supplemental Table 1** and comparisons to other reports in the literature are shown in **Supplemental Table 2**.

**Figure 1:**
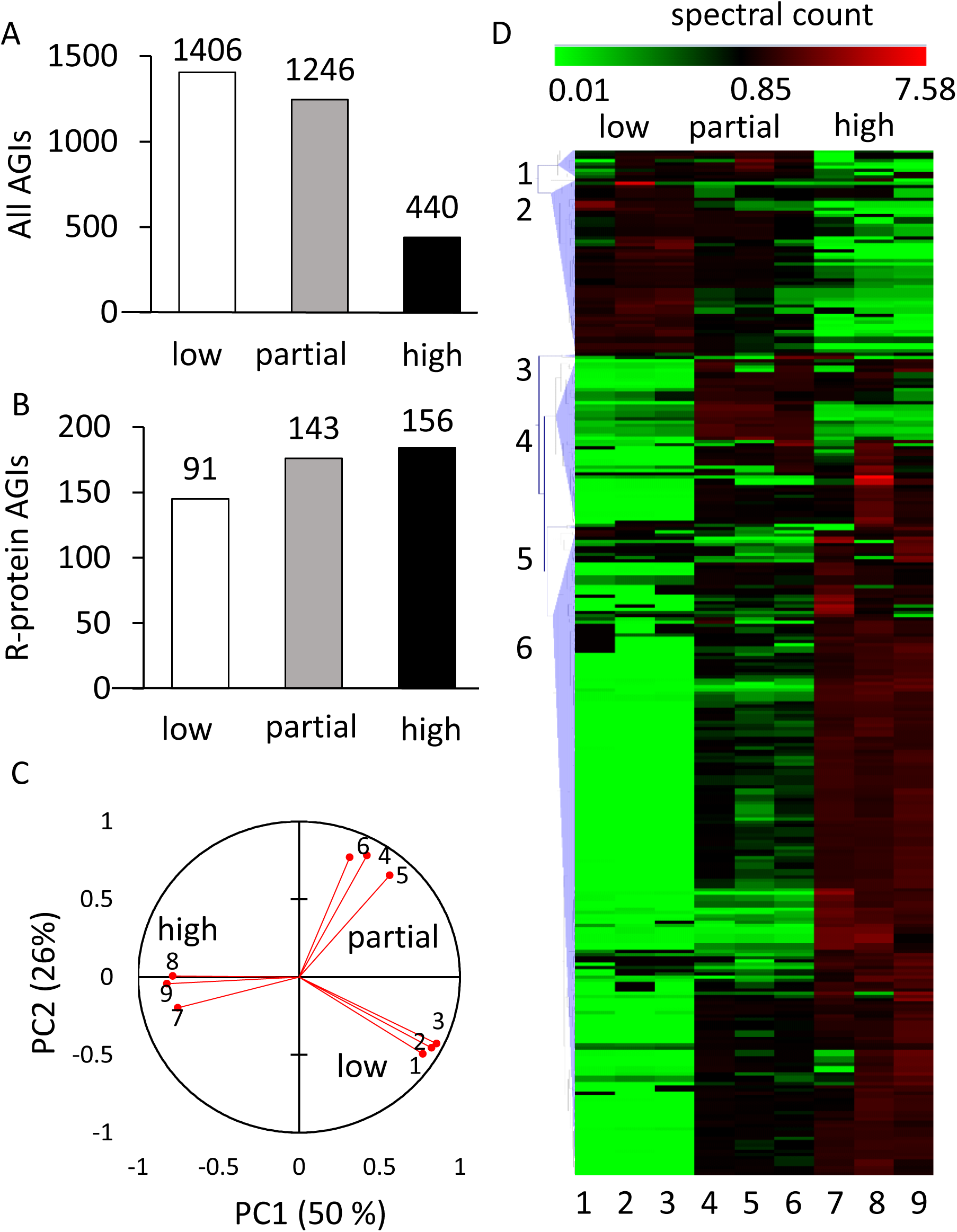
Purification and composition of the 80S cytosolic ribosome in Arabidopsis thaliana cell culture. A. Number of identified Arabidopsis proteins (AGIs) found in all 3 replicates following differential centrifugation steps for low, partial and high enrichment of ribosomes. B. The number of Arabidopsis 80S r-proteins isoforms identified across the three replicates. C. Principal component analysis of spectral counts assigned to the 319 proteins found by the differential centrifugation process across the three replicates and three purification steps (1-9). D. Hierarchical clustering analysis of spectral counts for the 319 proteins identified in the purification process. The green colour represents low and red colour high relative numbers of normalised spectral counts. The vertical numbers (1-6) indicate cluster number based on Pearson correlation. Cluster 6 contains 193 proteins, including 135 80S cytosolic ribosome r-proteins. The horizontal numbers (1-9) indicate replicate number as in C.

The intensity information for each peptide was extracted from the raw MS data files to estimate their relative abundance across the three ribosome enrichment steps. A total of 6084 peptides detected in all nine samples (three enrichment steps, three biological replicates) were included in the analysis **(Supplemental Table 3)**. Where more than one peptide per protein was available, a protein intensity value was then calculated by averaging the peptide values on a protein basis per replicate. This analysis resulted in quantitation of enrichment for 319 non-redundant proteins derived from ribosome preparations. PCA analysis of the peptide quantification data showed that two principal components explained more than 75% of the variation **(Figure 1C)**. The clustering of samples in the PCA demonstrated that the samples analysed were distinct between enrichment steps but consistent within biological replicates.

The protein intensity values were analysed by heirachical clustering and Pearson correlation to assess the similarity of the enrichment profiles of known r-proteins with non-ribosomal proteins in the set of 319 proteins. Six clusters based on a minimum similarity of 0.70 (Lamesch et al., 2012) were chosen for further analysis. Cluster 6 was the largest cluster and contained 193 proteins **(Figure 1D)**, including 135 cytosolic r-proteins. This cluster was significantly enriched in cytosolic r-proteins (p≤ 0.00001 Fisher’s exact) when compared to the whole proteome. Functional biological annotation of these 193 proteins showed that more than 70% (141 proteins) were annotated as components of protein synthesis and 135 of the 141 were annotated as located in the cytosol (based on SUBACon (Hooper et al., 2017); http://suba.live/); the remaining proteins were annotated to be located in the plastid or mitochondrion **(Supplemental Figure 1)**. These results indicated the methodology employed successfully enriched 80S cytosolic r-proteins. A total of 11 putative r-protein isoforms were identified that have not been previously reported in *Arabidopsis thaliana* ribosome purifications **(Table 1, Supplemental Table 2 for details).**

**Table 1:** Newly identified and re-annotated *Arabidopsis thaliana* cytosolic ribosomal protein genes. The proteins that shown here were not presented in the published Arabidopsis ribosome proteome data (Chang et al., 2005; Giavalisco et al., 2005; Carroll et al., 2008; Hummel et al., 2015).

| Family | AGIs | Protein name | Number of unique peptides identified |
| --- | --- | --- | --- |
| L18a | At1g29965 | RPL18aD | 9 |
| L24 | At2g36620 | RPL24A | 7 |
|  | At2g44860 | RPL24C | 3 |
| L36 | At2g37600 | RPL36A | 5 |
| L37 | At1g15250 | RPL37A | 2 |
| P2 | At3g44590 | RPP2D | 3 |
|  | At5g40040 | RPP2E | 9 |
| S15 | At5g43640 | RPS15E | 2 |
| S21 | At5g27700 | RPS21C | 2 |
| S27a | At1g23410 | RPS27aA | 5 |
|  | At2g47110 | RPS27aB | 5 |

### Enrichment of ribosome-associated proteins

To determine if the r-protein set and the other ribosome-associated protein set in Cluster 6 differed significantly in their enrichment, or if they came from the same distribution, the ratio of abundances (partial/low enrichment), (high/partial enrichment), and (high/low enrichment) were calculated for each protein in both groups. The results showed a non-normal distribution between these two sets with the value of the D statistic being 0.75, 0.40 and 0.66, respectively for each enrichment comparison (**Figure 2A**). The corresponding p-value using the Kolmogorov-Smirnov test suggests a significant difference between the enrichment characteristics of the r-protein and ribosome-associated protein sets (p < 0.0001). The distributions of these enrichments as histograms of the two protein sets are shown in **Figure 2B**. Even though the enrichment characteristics were different, it is possible that the 58 ribosome-associated proteins are part of a less tightly-bound translatome complex that partially dissociates during purification. As a further stringency filter the high/low enrichment of non-ribosomal proteins was set at a minimum of 10-fold with a p-value ≤ 0.05. Twenty-six of the 58 proteins met this enrichment criteria and are shown in **Table 2** as significantly enriched ribosome-associated proteins.

**Table 2:** Non-ribosomal proteins found to be ribosome-associated proteins in Arabidopsis cell culture. Non-ribosomal proteins were selected based of their >10-fold enrichment in the third centrifugation compared to the first centrifugation (p ≤ 0.05). The protein description (TAIR10), and significant fold enrichment (high/low) is shown.

| AGIs | Fold Enrichment (high/low) | P-value | Protein Description |
| --- | --- | --- | --- |
| AT5G53460.1 | 140 | 0.049 | NADH-dependent glutamate synthase 1. |
| AT5G20010.1 | 124 | 0.000 | RAS-related nuclear protein-1. |
| AT1G62850.2 | 120 | 0.041 | Class I peptide chain release factor. |
| AT5G55730.1 | 118 | 0.001 | Fasciclin-like arabinogalactan 1. |
| AT4G13780.1 | 83 | 0.045 | Methionyl-tRNA synthetase. |
| AT3G09840.1 | 63 | 0.043 | Cell division cycle 48. |
| AT3G53230.1 | 63 | 0.050 | ATPase, AAA-type, CDC48 protein. |
| AT3G27850.1 | 59 | 0.001 | 50S ribosomal protein L12-C. |
| AT1G48920.1 | 55 | 0.020 | Nucleolin like 1. |
| AT2G27030.3 | 53 | 0.041 | Calmodulin 5. |
| AT2G01140.1 | 50 | 0.042 | Aldolase superfamily protein. |
| AT2G21330.1 | 50 | 0.049 | Fructose-bisphosphate aldolase 1. |
| AT1G22840.1 | 48 | 0.000 | Cytochrome C-1;encodes cytochrome c. |
| AT3G09820.1 | 48 | 0.010 | Adenosine kinase 1. |
| AT5G03300.1 | 48 | 0.010 | Adenosine kinase 2. |
| AT4G05400.1 | 33 | 0.000 | Aopper ion binding. |
| AT1G08580.1 | 31 | 0.004 | Not assigned. |
| AT4G35460.1 | 28 | 0.006 | NADPH-dependent thioredoxin reductase B. |
| AT2G17420.1 | 28 | 0.026 | NADPH-dependent thioredoxin reductase A. |
| AT2G21870.1 | 27 | 0.049 | Copper ion binding;cobalt ion binding;zinc ion binding. |
| AT3G51160.1 | 26 | 0.001 | NAD(P)-binding Rossmann-fold superfamily protein. |
| AT3G58610.1 | 26 | 0.048 | Ketol-acid reductoisomerase; |
| AT3G55620.1 | 15 | 0.038 | Translation initiation factor IF6. |
| AT1G15270.1 | 15 | 0.004 | Translation machinery associated TMA7. |
| AT3G55410.1 | 13 | 0.017 | 2-oxoglutarate dehydrogenase. |
| AT5G08570.1 | 10 | 0.039 | Pyruvate kinase family protein. |

**Figure 2:**
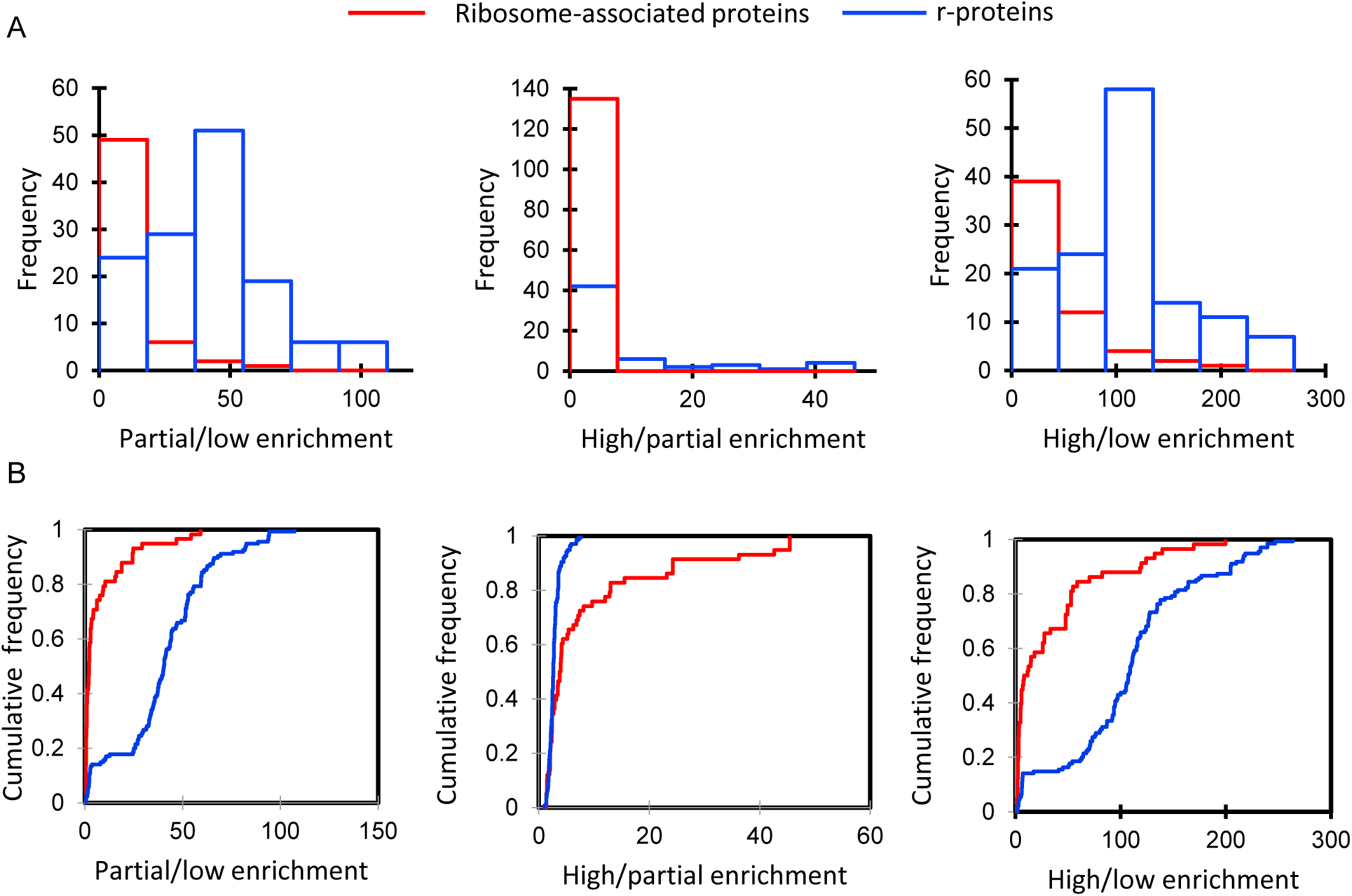
Frequency distribution graphs of the ratio of spectral counts for r-proteins and ribosome-associated proteins in pairs of ribosomal enrichments. A. Histogram of the ratio of enrichment based on spectral counts for r-proteins and ribosome-associated proteins; blue colour indicates the distribution of r-proteins and red colour indicates the distribution of ribosome-associated proteins. B. Cumulative relative frequency distribution graphs; statistical significance was estimated by the Kolmogorov-Smirnov test for 193 proteins including 135 r-proteins (blue) and 58 ribosome-associated proteins (red) (p < 0.0001) from Cluster 6 (Figure 1D).

### Protein turnover rates of ribosomal proteins

As a further criteria to verify if proteins were part of a ribosomal complex *in vivo*, we sought to calculate the protein turnover rate of proteins in ribosomal extracts in *Arabidopsis thaliana* cell culture to determine which proteins turned over as a similar rate. We had previously shown the distribution of turnover rates in *Arabidopsis thaliana* cell culture varied more than 100-fold (Li et al., 2012) but that members of the same complex tend to turnover at a similar rate (Li et al., 2013). Cell cultures were labelled by changing the growth medium from a solution that contained inorganic ^14^N to one that only contained inorganic ^15^N. As proteins degrade over time, newly synthesized proteins increased in the amount of ^15^N they contained as the amino acid pool was progressively labelled. Calculation of the fraction of the protein labelled with ^15^N (labelled peptide fraction or LPF) could be achieved by a binary differentiation between the natural abundance population before labelling and the newly synthesised labelled population of peptides by a non-linear least squares analysis (Li et al., 2017). Determining the proportional degradation rate (K_D_) and a normalised synthesis rate (Ks/A) for each protein requires the LPFs, the growth rate of the cell culture calculated by determining the fresh weight (g) of the cell culture from day 1 to day 5 **(Figure 3A)**, and a series of calculations as previously reported (Li et al., 2017).

**Figure 3:**
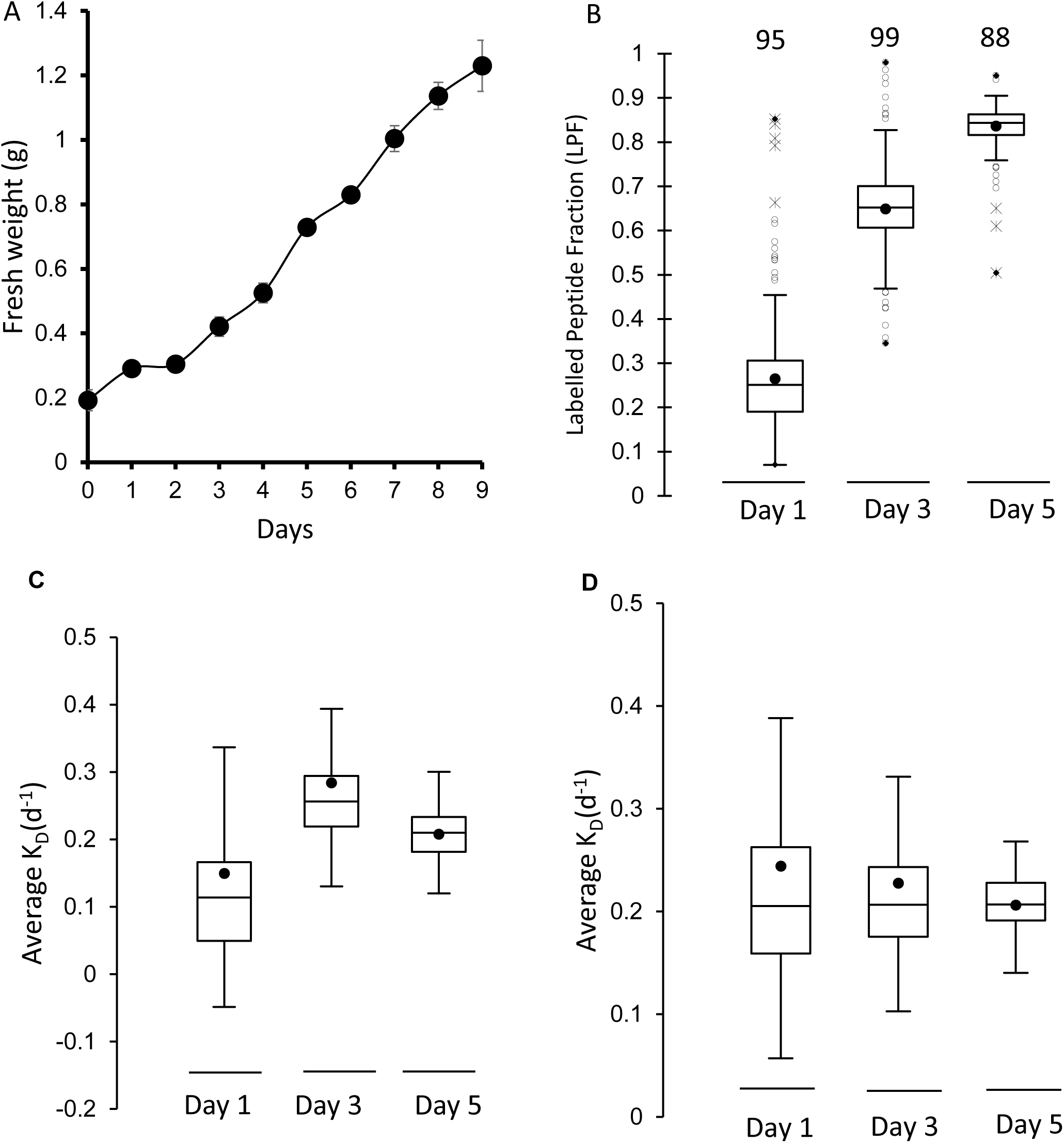
Protein turnover of ribosomes in *Arabidopsis thaliana* cell culture. A. Growth rate of *Arabidopsis thaliana* cell culture over 9 days after transferred into new media (FW, n=4, standard error). B. Box plot of the average ^15^N enrichment of r-protein peptides after 1, 3 and 5 days of growth in 98% ^15^N media. The number of r-proteins included for each time point is shown above the box plot (n=4). C,D box plots representing the average degradation rate (K_d_) of r-proteins after day 1, 3 and 5 through the course of the growth experiment. C. K_d_ for each time point using measured FCP values. D. K_D_ using the median polish strategy for FCP calculation.

LPFs of ribosomal protein in *Arabidopsis thaliana* cell culture were calculated from three times points during the growth of the cell culture (day 1, 3 and 5) using four biological replicates. Highly enriched ribosomal protein fractions from the cytosol was isolated as indicated above, digested by trypsin and the peptides analyzed by LC-MS/MS. The ^15^N enrichment of the labelled peptide fractions were calculated at each time point, showing ^15^N enrichment in new peptides was ∼ 60% after 1 day and rose to 83% by day 5 **(Supplemental Figure 2)**. Calculated LPF showed a clear increase from 25% at 1 day to more than 85% by day 5 **(Figure 3B)**. From LPF data the degradation rate for protein isoforms were calculated at each timepoint with the criteria that peptides for a protein were found in three or more biological replicates and the protein calculation was supported by at least five independently quantified peptides. The measured K_D_ of these proteins over day 1, day 3 and 5 showed variability based on fluctuations in FCP value coming from measuring the fresh weight of *Arabidopsis thaliana* cell culture **(Figure 3C)**. To gain a more precise degradation rate in each replicate a median polish strategy (Li et al., 2017) was used for experimental normalization and to calculate FCP (Figure 3D). The overall correlation between the measured FCP and calculated FCP was very high (R = 0.96), and the median polish approach gave a more precise estimate of growth of the cell culture in each labelled replicate, lowering K_D_ variation.

Of the proteins in the cytosolic ribosome of *Arabidopsis thaliana*, the degradation rate (K_D_) and relative synthesis rate (K_s_/A) from 54 protein families encoded by 89 genes were calculated. These proteins were all r-proteins rather than the ribosomal-associated proteins; presumably due to the lower abundance of the latter. The turnover rate set included proteins from 29 r-protein families from the large subunit **(Figure 4A)** and from 25 r-protein families from in the small subunit **(Figure 4B)** which were encoded by 49 and 40 genes, respectively.

**Figure 4:**
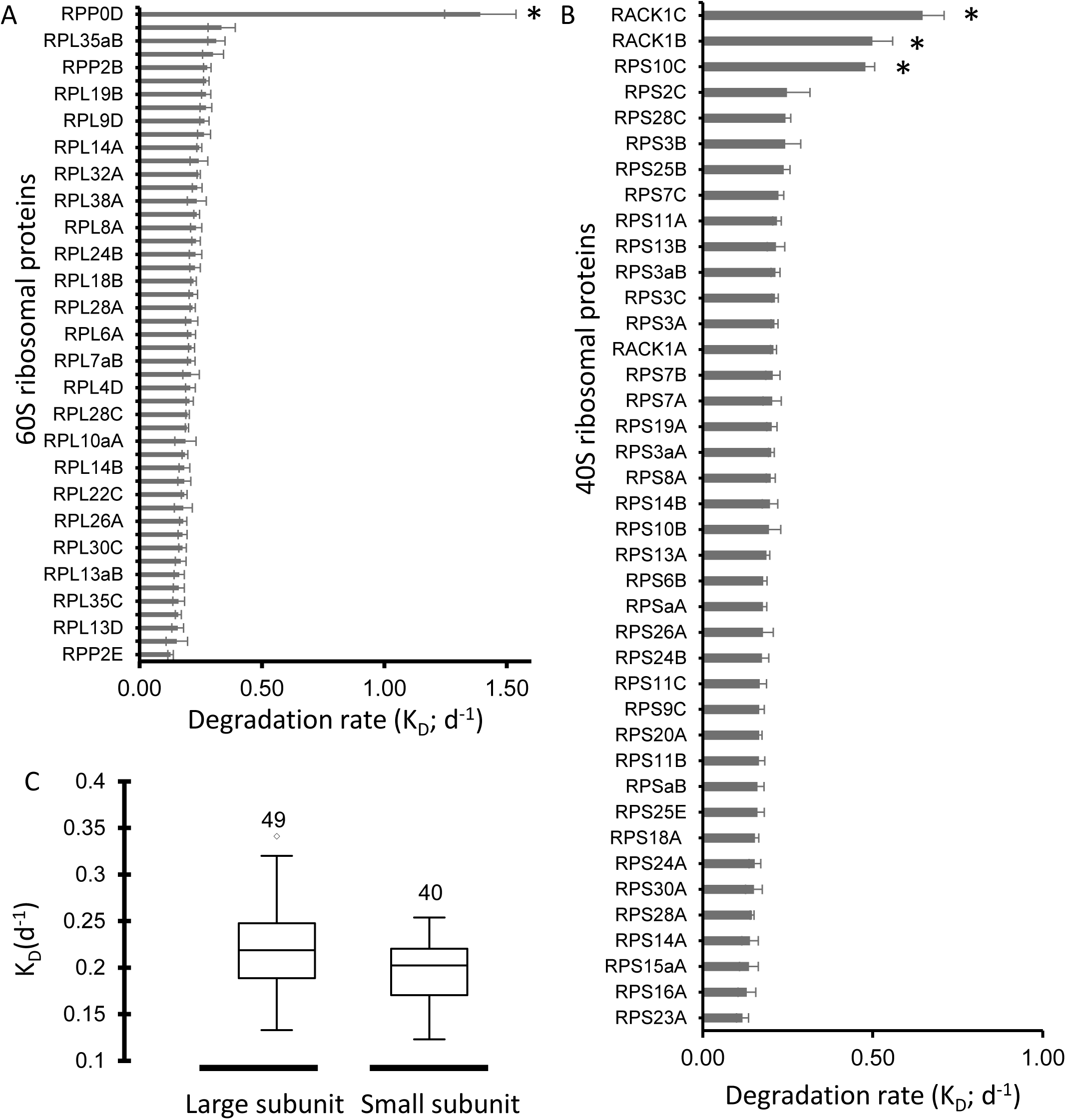
Turnover rate of specific r-proteins in Arabidopsis cell culture. A. Degradation rate of 60S ribosomal subunit proteins. B. Degradation rate (KD) of 40S ribosomal subunit proteins. In both A and B the means for each protein was calculated from KD data for ≥ 3 biological replicates and quantified from ≥ 5 peptide spectra. Error bars: standard error. * indicates significant difference from the mean of the ribosome (based on T-Tests p<0.05). C. Box plots of K_D_ values for r-proteins of the large and small subunits.

More than 90% of the r-protein degradation rates fell within a narrow range of 0.18-0.25 d^-1^ (half-lives of 3-5 days) **(Supplemental Table 4).** There was no significant difference in degradation rate between 60S ribosomal subunits and 40S subunits with averages of 0.25 and 0.22 d^-1^, respectively **(Figure 4C)**. Relative synthesis rate (K_s_/A) data mirrored the degradation rate data for each r-protein isoform **(Supplemental Table 4).** However, there were several r-proteins that showed degradation and synthesis rate characteristics that were significantly different to those of the other r-proteins in each ribosome subunit. In the 60S ribosomal subunit, the r-protein RPP0D (60S acidic ribosomal protein family/ribosomal protein L10 family) was very unstable with a K_D_ of 1.39 d^-1^ and Ks/A of 1.7 d^-1^ and thus a half-life of 0.5 d. In the 40S ribosomal subunit, two isoforms of RACK1, namely RACK1b and RACK1c, had K_D_ of 0.5-0.65 d^-1^ and Ks/A of 0.74-0.87 d^-1^ and half-lives of 1-1.4 days. Indepth visual inspection the LPF data supporting these exceptional degradation and synthesis rates **(e.g. Supplemental Figure 4)** confirmed the K_D_ and Ks/A calculations.

## Discussion

### Core 80S ribosomal r-protein composition in plants defined by quantitative enrichment, co-synthesis and co-degradation

This analysis has provide the most robust foundation yet for the Arabidopsis r-protein content of 80S ribosomes by providing multiple lines of evidence to justify protein inclusion in the core r-protein set. We now have information on the degree of enrichment during ribosome purification and the *in vivo* turnover rate of putative r-proteins in Arabidopsis **(Supplemental Table 3)**. A full numerical comparison showing the improvement in coverage over previous research (Chang et al., 2005; Giavalisco et al., 2005; Carroll et al., 2008; Hummel et al., 2015) is also shown graphically in **Supplemental Figure 4A,B.**

Five of the r-proteins first identified here (Table 1) are proposed by homology to be located in the 60S subunit and six are proposed to be in the 40S subunit (Chang et al., 2005; Giavalisco et al., 2005; Carroll et al., 2008; Hummel et al., 2015). In total at least one isoform of 76 out of 81 ribosomal protein families predicted in Arabidopsis were identified in this analysis. This included 108 protein isoforms in the 60S large subunit and 76 protein isoforms in the 40S small subunit **(Supplemental Table 3)**.

Comparing the r-proteins observed here and in previous reports with a systematic assessment of r-protein families and groups from other kingdoms of life, did not reveal any significant difference between the turnover rate or fold-enrichment during ribosome purification for eukaryotic specific subunits versus those present across all kingdoms **(Supplementary Table 4)**. The eukaryotic protein families predicted in plants but not found experimentally to date in Arabidopsis are the large subunit r-proteins RPL29 (RPL29A and RPL29B), RPL39 (RPL39A, RPL39B and RPL39C), RPL40 (RPL40A and RPL40B) and RPL41 (RPL41C, RPL41D, RPL41E and RPL41G). This is likely because their short amino acid sequences (25-128 AAs) and many Arg and Lys residues yield short and few tryptic peptides which are not easily detected by mass spectrometry (Hummel et al., 2015).

### Turnover of ribosomal proteins

In whole tissue studies in yeast (Christiano et al., 2014), Barley (Nelson et al., 2014) and Arabidopsis (Li et al., 2017) and it had been shown that turnover rates of r-proteins fall within a relatively narrow range and are among the longest-lived protein complexes in cells. We have confirmed this is also the case for isolated Arabidopsis ribosomes and that degradation rates and synthesis rates of proteins in both 60S and 40S subunits were very similar **(Figure 4C, Supplemental Table 4**. However, we did observe three bonefide Arabidopsis r-proteins with unusually rapid degradation and synthesis rates, RACK1b, RACK1c and RPP0D, respectively.

RACK1b and RACK1c have been previously found in ribosomes (Carroll et al., 2008; Hummel et al., 2015) and their function as r-proteins subunits has been investigated (Guo and Chen, 2008; Guo et al., 2009). RACK1 is required for efficient translation of mRNA during the translation initiation process within the 40S ribosomal subunit (Thompson et al., 2016). RACK1 also acts as a scaffold protein between ribosomal machinery and many signalling pathways in the cell (Islas-Flores et al., 2015). In *Arabidopsis*, knockout of RACK1A results in developmental defects (Chen et al., 2006) and hypersensitivity to hormonal treatment (Guo et al., 2009). Knockout of RACK1B or RACK1C alone did not lead to significant phenotypes in the presence of RACK1A, but their loss exacerbated the *rack1a* phenotype in double mutants leading to the suggestion they are functionally redundant with RACK1a (Guo and Chen, 2008). A more specialised role for RACK1B,C might be infered from the combination of these data with our evidence of rapid turnover rates for RACK1B and RACK1C, but a typical slow turnover rate for RACK1A.

RPP0D was previously reported as a ribosomal subunit isolated from Arabidopsis cell culture (Carroll et al., 2008), however it was not reported in ribosomal purifications from Arabidopsis leaves or whole seedlings (Chang et al., 2005; Giavalisco et al., 2005; Hummel et al., 2015). Examination of whole Arabidopsis proteome datasets from multiple tissues show the RPP0D protein is present in cell culture, flowers and siliques, but not in vegetative tissues (Baerenfaller et al., 2008). Carroll et al. (2008) named this protein RPP0D based on its presence in ribosomes and its homology with other P0 isoforms. But the homology with other RPP0 sequences is only in one region of the protein and the overall sequence identity is only 20%. Re-analysis by sequence homology searches show that RPP0D shares more than 60% sequence identity to a yeast and human protein called Mrt4. Mrt4 is not P0 isoform but rather it is a pre-60S trans-acting factor (Kemmler et al., 2009; Lo et al., 2009; Rodriguez-Mateos et al., 2009). Mrt4 shares sequence similarity to P0 proteins in yeast and humans, and P0 together with P1/P2 and L12 form the ribosomal stalk responsible for translation factor-dependent GTP hydrolysis (Michalec et al., 2010). Mrt4 binds to pre-60S during its shuttling between the nucleus and the cytoplasm at the same site in the ribosome as P0 but is displaced by P0 in the cytosol on ribosome maturation (Michalec-Wawiorka et al., 2015). Our data suggest that the very rapid degradation rate of RPP0 in Arabidopsis and its homology with Mrt4 warrents the renaming of At1g25260 as Mrt4 and its further investigation in ribosome biogeneis as a pre-60S trans-acting factor.

### Ribosome-associated proteins as potential translatome components in Arabidopsis

Identification and further characterization of ribosome-associated proteins that are distinct from r-protein families are essential in order to obtain a better understanding of the physical organisation of the process of translation and post-translational folding and modification of nascent polypeptides in plants. An earlier study in yeast showed that proteins that are associated with the ribosome are enriched in proteins needed for the translational process; so called translation-machinery-associated (TMA) proteins. Deletion of some of these proteins (TMA7) resulted in an alteration in both protein synthesis rate and translation-related processes in yeast (Fleischer et al., 2006).

Here we identified a significant number of ribosome-associated proteins in the purification process of ribosomes in Arabidopsis. By using the open source STRING database version 10.5 (https://string-db.org/) (Szklarczyk et al., 2015) the protein interaction score was compared between known r-proteins in Arabidopsis and these ribosome-associated proteins. Most of the proteins that are considered ribosome-associated proteins in this study showed a significant interaction score with r-proteins based on a combination of co-expression and protein-protein interaction data, confirming their likely associations with ribosomes **(Supplemental Table 3)**. By imposing a strict definition of enrichment on our data of ≥ 10 fold enrichment during co-purification, 26 putative ribosome-associated proteins are proposed for Arabidopsis (**Table 2**).

Eight of these proteins have been previously found to be involved in translation and ribosomal function in different organisms, including TMA7 (Fleischer et al., 2006), eIF6 (Guo et al., 2011), cytochrome *c* (Mei et al., 2010), Class I peptide chain release factor (Petropoulos et al., 2014), methionyl-tRNA synthetase (Kwon et al., 2011), nucleolin like 1 or PARL1 (Petricka and Nelson, 2007), fructose-bisphosphate aldolase 1(Ziveri et al., 2017) and AAA-ATPases (Bassler et al., 2010). AT1G15270, which is annotated as Arabidopsis gene translation machinery associated TMA7, gained this annotation based on homology to the yeast protein (Fleischer et al., 2006). However, to our knowledge this is the first direct experimental evidence of its association with ribosomes in a plant species. No further information on its function in yeast has yet been revealed since the Flesicher et al. (2006) report. AT3G55620, annotated as a translation initiation factor IF6, is also shown to be ribosome-associated in our study. This protein has been found in Arabidopsis to physically interact with the small ribosomal RACK1 protein (Guo et al., 2011) as well as to be involved in the formation of the 60S large subunit (Brina et al., 2015). eIF6 in yeast (*S.cerevisiae*) is required for pre-rRNA processing and the biogenesis of 60S ribosome subunit, and the depletion of this protein resulted in imbalance of the 60S to 40S subunit ratio and reduction of protein synthesis and eventually reduction of cell growth (Basu et al., 2001). Unexpectedly, cytochrome *c*, a mitochondrial protein involved in electron transport, was found in the ribosomal enrichment list (Table 2). In mammals this protein has been reported to be a tRNA-associated protein that is essential for disabling the formation of an apoptosis complex in the cytosol (Mei et al., 2010). It is still unclear the functionality of other proteins that interacts with the 80S cytosolic ribosome and how they participate, if at all, in protein synthesis in plant cells. Further study is required to reveal the role of these ribosome-associated proteins during the translational process in plant cells.

## Methods

### Arabidopsis thaliana cell culture

*Arabidopsis thaliana* cell suspension (ecotype Landsberg erecta) was cultured in growth medium 1x Murashige and Skoog (MS) Modified Basal Salt Mixture (phyto technology M524) without vitamins, 3% w/v sucrose (Ajax chemical A530), 0.5 mg/L naphthaleneacetic acid (Sigma Aldrich 15165-79-4), 0.05 mg/L kinetin (Sigma Aldrich K0753), pH 5.8 at 22°C with orbital shaking at 100-120 rpm under constant light (100 µmolm^-2^s^-1^). Cultures were maintained in 250 ml Erlenmeyer flasks by the inoculation of 20 ml of 7 days old cells into 100 ml fresh growth medium.

### ^15^N labelling of Arabidopsis thaliana cell culture

Seven day old *Arabidopsis thaliana* cell culture was transferred from non-labelled media (^14^N) to media without nitrogen, and the cells washed three times to eliminate all ^14^N in the media. Washed cells were transferred to heavy media (^15^N) containing two nitrogen sources for optimal growth (Li et al., 2012) including 1.65 g/L ^15^NH_4_^15^NO_3_ (Sigma Aldrich 299278) and 1.9 g/L K^15^NO_3_ (98% ^15^N Sigma) (Sigma Aldrich 335134). To study protein turnover rate, the labelled *Arabidopsis thaliana* cell culture was collected by vacuum filtration after the first day, third day and fifth day of transformation into new media, then stored in a -80°C freezer until use.

### Ribosome purification from cell culture

To isolate 80S cytosolic ribosomes from Arabidopsis cell culture, 10 g of 7 day old *Arabidopsis thaliana* cells were frozen in liquid nitrogen and ground in a pre-cooled mortar with a pestle. The powder was transferred into extraction buffer containing 0.45 M mannitol (Ajaxs chemical A310), 30 mM HEPES (Sigma Aldrich H3375), 100 mM KCl (Sigma Aldrich P9333), 20 mM MgCl_2_.6H_2_O (Ajax Finechem A296), 0.5% (w/v) polyvinyl pyrrolidone-40 (Sigma Aldrich 9003-39-8) and 0.5% (w/v) bovine serum albumin (Bovogen biologicals BSAS1.0) with addition of 20 mM cysteine-L (Sigma Aldrich A-9165) prior to grinding and pH adjusted to 7.5, followed by using an ultra-turax for homogenization of the mixture. The homogenised solution was filtered through two layers of miracloth (Merk Millipore, Darmstadt, Germany). The filtrate was transferred into a pre-cooled, sterilized 25 ml of Beckman Coulter centrifuge tubes and centrifuged at 30000 *xg* for 20 minutes at 4°C. The supernatant (low purity ribosome extract) was transferred into 1.5 M sucrose (Ajaxs chemical A530) dissolved in a mixture of 2x ribosome resuspension buffer consisting of 30 mM HEPES, 100 mM KCl and 20 mM MgCl_2_.6H_2_O, pH adjusted to 7.5 and centrifuged at 60000 rpm for 90 minutes. After the first ultra-centrifugation, the pellet (partially purified ribosome extract) was resuspended in 2 ml of a solution that consisted of 5 mM DTT (Sigma Aldrich D0632) and 1.5 M sucrose. The resuspended sample was mixed with 2x ribosome extraction buffer followed by second ultra-centrifugation at 60000 rpm for 2.5 hours. Finally, the purified ribosome pellet was resuspended in 1x ribosome extraction solution and stored in -80 °C for later use. Ribosomal protein content was quantified using Amido black against a standard solution of bovine serum albumin.

### Precipitation, denaturation and digestion of ribosomal proteins

Ribosomal extracts (50 µg protein) were precipitated in cold acetone and incubated for 1 hour in - 80 °C and centrifuged for 20 minutes at 20800 *xg* at 4°C. The protein pellet was denatured, reduced and alkylated using 1.5 M urea, 10 mM DTT and 25 mM iodoacetamide. Denaturated protein was digested using trypsin (Invitrogen MSI0015) and incubated overnight at 37°C and the reaction terminated by acidification of the mixture with 1% formic acid. Peptides from digested protein was purified and concentrated before mass spectrometry analysis using micro spin column silica C18 in a reverse phase column.

### LC-MS/MS and Data analysis

Mass spectrometry analysis was performed with an Thermo Scientific Orbitrap Fusion mass spectrometer. The raw data files (. raw) files were converted to .mzML files by using open source MS convert software. Then the .mzML files were converted to .mgf files via the Convert mZ[X]ML tool in the trans-proteomic pipeline v. 4.8 (TPP). The Mascot search algorithm (Matrix Science) was used to search tandem mass spectra against proteins from the TAIR 10 release of *Arabidopsis thaliana* (https://www.arabidopsis.org/) database to convert the. mgf files to both .dat and .csv files. Search parameters were: variable modifications: carbamidomethyl (C) and oxidation (M), monoisotopic, trypsin selected as digesting enzyme, and maximum missed cleavages set to one. Moreover, the peptide charges chosen as (+2, +3 and +4), ^13^C error set to one and peptide tolerance selected as ± 50 ppm. Finally, MS/MS ion searches were activated, and MS/MS tolerance selected as ± 0.6 Da. Mascot results were exported as a .dat file, and then converted to pep.xml using the PepXML tool in TPP for determination of peptide and protein probabilities. Pep.XML files were then further converted to interact.pep.XML by using the peptide prophet tool in TPP. Peptide level LPFs were estimated from the interact.pep.xml and the mzML files by an in-house solution as described previously (Nelson et al., 2014), rewritten in R and including an extra filter. The heavy population, calculated by substracting the natural abundance envelope from the observed isotopes as described in (Nelson et al., 2014), is now also filtered by calculating the correlation coefficient between the estimated heavy distribution as per NNLS result and the observed data. We found this correlation filter to more accurately remove data with poor signal-to-noise or contamination by other peptides than the fit of a gaussian distribution. The minimum correlation to pass the filter was set to 0.5. 15 seconds was chosen as retention time tolerance, m/z tolerance was 10 ppm, and 95% selected as maximum label enrichment. For ^15^N labelled protein studies nitrogen (N) was chosen as the labelled element and 15 as the isotope number. For each sample that was analyzed by this pipeline a .tsv file was generated. Protein level LPFs were obtained from these .tsv files in conjunction with TPP .prot.xml files by taking the median of peptide LPFs for each protein. The mass spectrometry proteomics data have been deposited to the ProteomeXchange Consortium via the PRIDE partner repository with the dataset identifier PXD012839. MS1 quantiation was performed using the Skyline MS1 filtering workflow. Peak integrations were manually inspected and areas of monoisotopic peaks exported for statistical analysis (t-test).

### Bioinformatics analysis

To check if the non-core ribosomal proteins identified in this study were related with core ribosomal proteins in previous studies we investigated seven score summaries from Arabidopsis String-db (https://string-db.org/cgi/download.pl) (Szklarczyk et al., 2015). This data scores the interaction between each pair of genes as summarised from transcript studies, protein:protein interaction studies etc. We calculated interaction scores for each of the genes whose protein product was identified in the final centrifugation by calculating an average of its interaction scores with a core set of bonafide ribosomal proteins as judged by their inclusion in previous Arabidopsis ribosomal studies.

### Statistical analysis

The analysis of data including one-way and two-way variance analysis (ANOVA) and Kolmogorov-Smirnov test for comparison of samples using XLSTAT version 2018.5, less than 0.05 was considered as significant. For ANOVA the criteria selected were: 0.0001 was chosen as tolerance, confidence interval was set at 95%. For model selection, we chose the best model with criterion (adjusted R2), and number 2 was selected as a minimum and maximum variables. For the Kolmogorov-Smirnov (K-S) test, the criteria were selected as follows: Alternative hypothesis F1(x) ≠ F2(x), 5% was chosen as significant level, hypothesised difference (D) = 0 and asymptotic p-value was selected.

## Supplementary Figures List

Supplemental figure 1: Functional categorisation of all proteins in cluster 6.

Supplemental Figure 2. ^15^N enrichment in r-protein peptides after different periods of time following transfer to ^15^N media.

Supplemental Figure 3: Histogram of the proportion of natural abundance and heavy labelled peptides for RPPOD and RACK1A,B and C peptides.

Supplemental Figure 4: Comparisons to number of r-proteins found in this study to previously published data.

## Supplementary Tables List

Supplemental Table 1: Number of total proteins and r-proteins identified in each of three replicates for each centrifugation purification step to produce low, partial and high purity ribosomal extracts from Arabidopsis cell culture.

Supplemental Table 2: Identification of Arabidopsis thaliana cytosolic r-protein genes and comparison to other published data

Supplemental Table 3: Arabidopsis thaliana cytosolic ribosome enrichment through differential centrifugation

Supplemental Table 4: Collated features of 80S cytosolic ribosome proteins in Arabidopsis thaliana including synthesis and degradation rates, r-protein families in other kingdoms and identification claims for r-proteins in other Arabidopsis reports.

## Primary Data

PRIDE Project Name: Ribosomal protein turnover under normal conditions Project accession: PXD012839

---

**Supplemental Figure 1:**
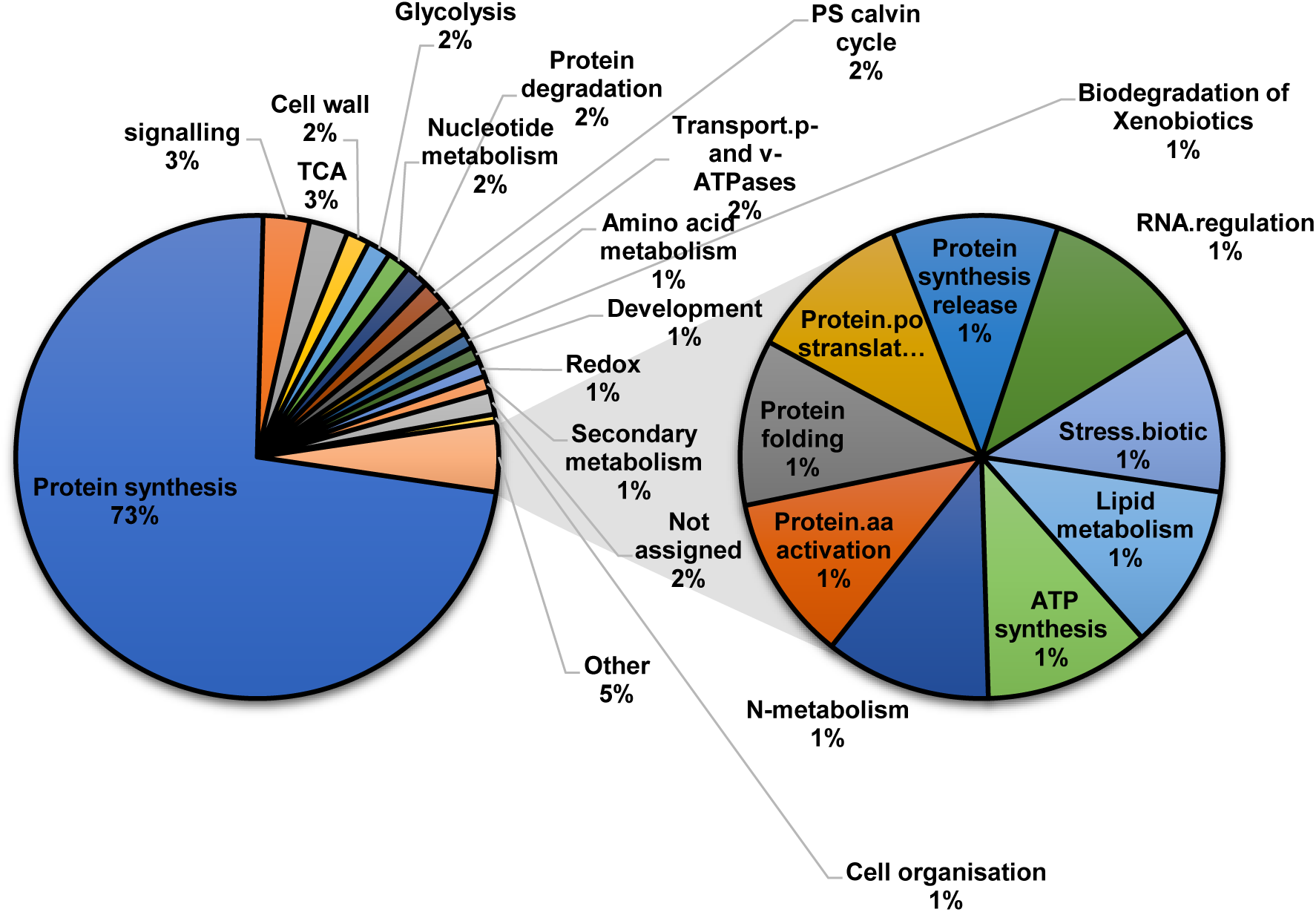
Functional categorisation of all proteins in cluster 6. Protein functional categories (MapMan) were assigned to the 193 proteins of cluster 6. 73% of the proteins (141 proteins) in cluster 6 are annotated as directly participating in protein synthesis while others are associated with preparations for protein synthesis and protein modification.

**Supplemental Figure 2.**
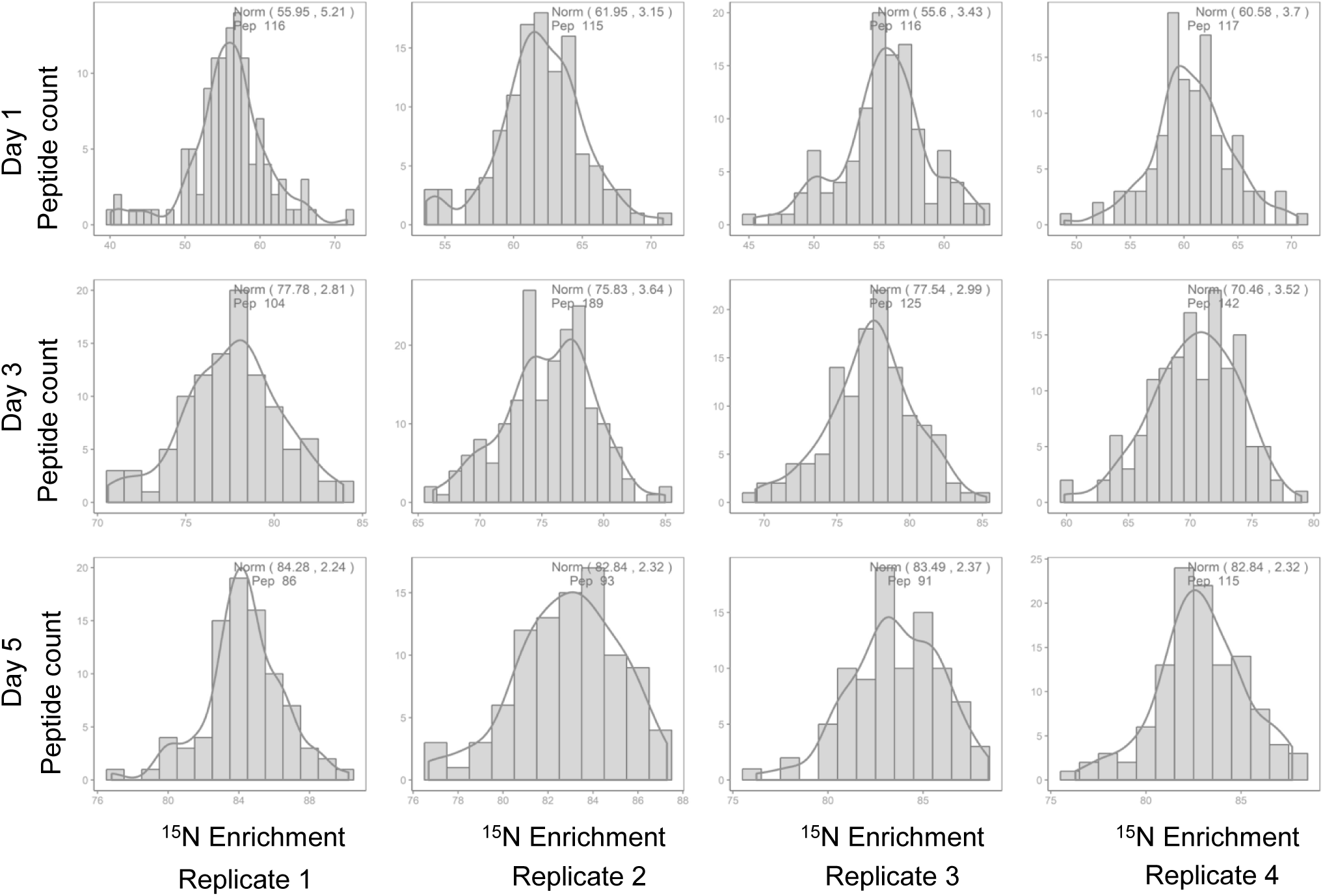
^15^N enrichment in r-protein peptides after different periods of time following transfer to ^15^N media. The grey bars are the actual peptide data, the median and standard deviation (x, y) and the number of peptides (pep) included in each analysis is shown. The plotted line is a normal distribution (norm).

**Supplemental Figure 3:**
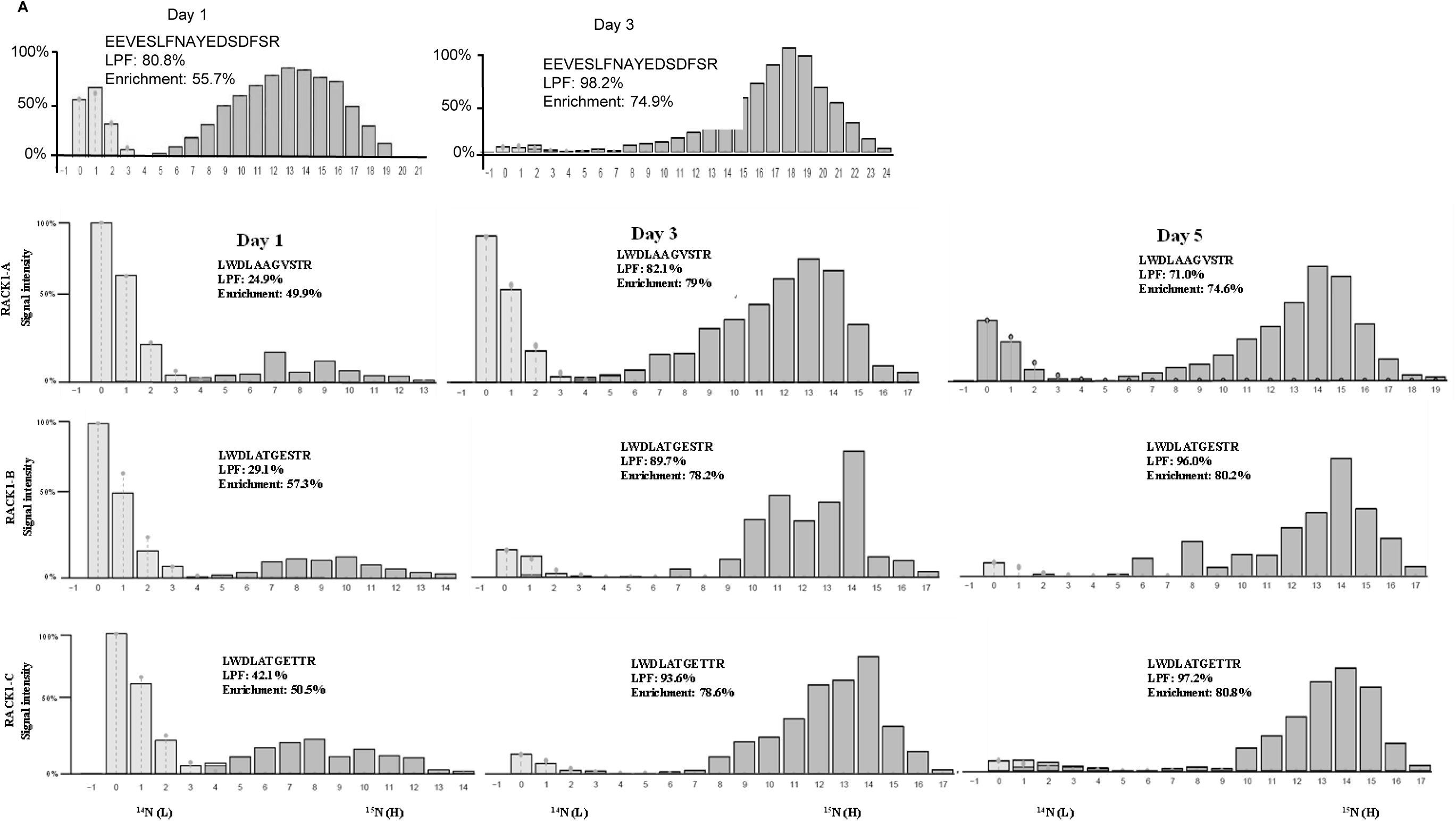
Examples of histograms of the proportion of natural abundance (light grey) and heavy labelled (dark grey) peptides used to calculate the labelled protein fraction (LPF) for RPPOD and RACK1A,B and C. (A) Labelling of RPPOD specific peptide EEVESLFNAYEDSDFSR at Day 1 and Day 3. (B) Labelling of specific peptides for RACK1A, B and C at Day 1,3 and 5.

**Supplemental Figure 4:**
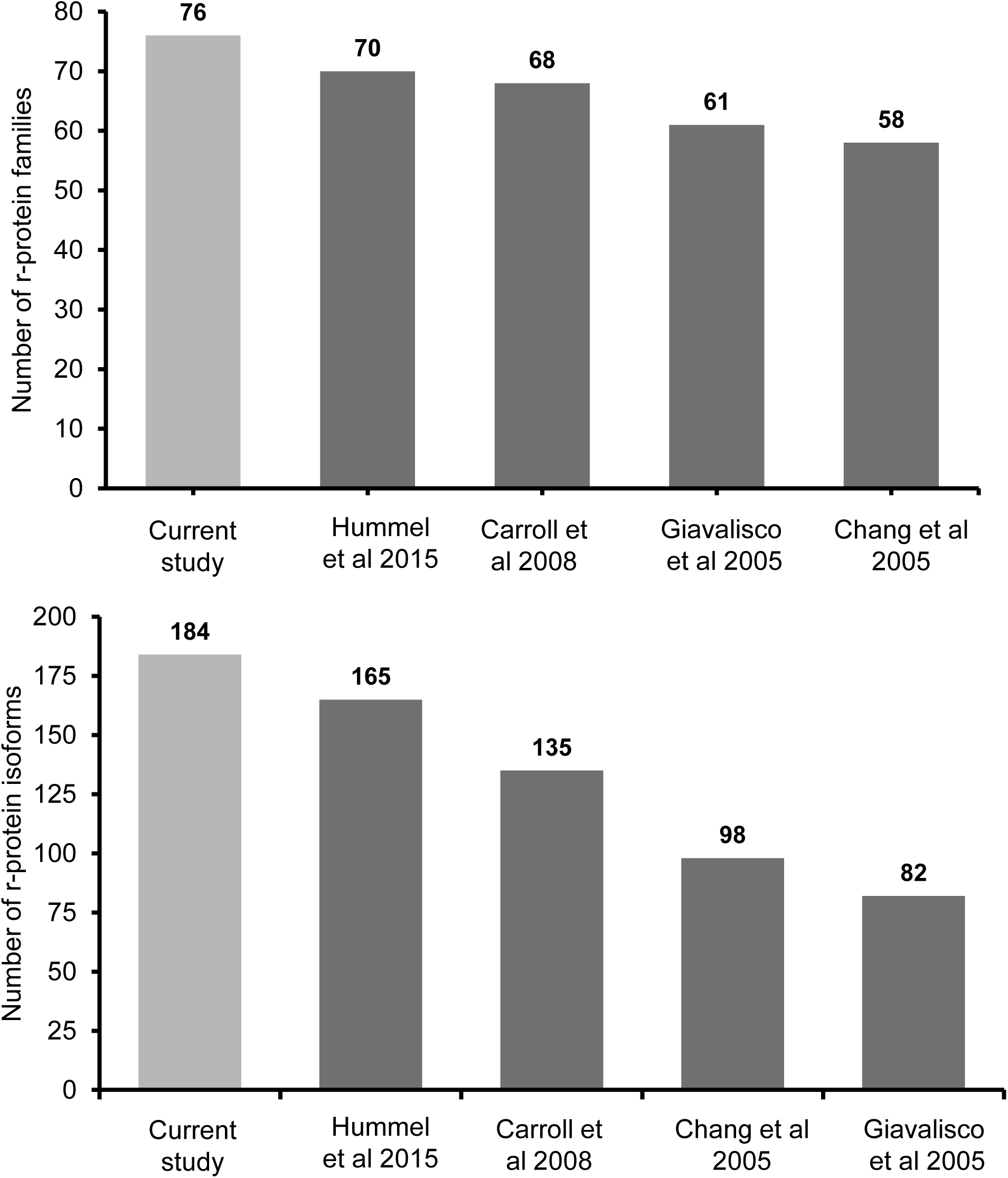
Comparison of number of r-protein families and r-protein isoforms found in this study (light grey) to previously published reports (dark grey). (A) Number of r-protein families that has been identified in comparison with findings of published reports. (B) Comparison of the number of r-protein isoforms identified in any of the analyses in the current study compared to previously published reports.

